# A missense variant impairing TRMT1 function in tRNA modification is linked to intellectual disability

**DOI:** 10.1101/817247

**Authors:** Kejia Zhang, Jenna M Lentini, Christopher T Prevost, Mais O Hashem, Fowzan S Alkuraya, Dragony Fu

## Abstract

The human *TRMT1* gene encodes a tRNA methyltransferase enzyme responsible for the formation of the dimethylguanosine (m2,2G) modification in cytoplasmic and mitochondrial tRNAs. Frameshift mutations in the *TRMT1* gene have been shown to cause autosomal-recessive intellectual disability (ID) in the human population but additional TRMT1 variants remain to be characterized. Moreover, the impact of ID-associated TRMT1 mutations on m2,2G levels in ID-affected patients is unknown. Here, we describe a homozygous missense variant in *TRMT1* in a patient displaying developmental delay, ID, and epilepsy. The missense variant changes a conserved arginine residue to a cysteine (R323C) within the methyltransferase domain of TRMT1 and is expected to perturb protein folding. Patient cells expressing the TRMT1-R323C variant exhibit a severe deficiency in m2,2G modifications within tRNAs, indicating that the mutation causes loss-of-function. Notably, the TRMT1 R323C mutant retains the ability to bind tRNA but is unable to rescue m2,2G formation in TRMT1-deficient human cells. Our results identify a pathogenic point mutation in TRMT1 that severely perturbs tRNA modification activity, and provide the first demonstration that m2,2G modifications are disrupted in patients with TRMT1-associated ID disorders.

## Introduction

The post-transcriptional modification of tRNA has emerged as a critical modulator of biological processes ranging from gene expression to development (Frye et al. 2018; Ranjan and Leidel 2019). There are over 100 types of tRNA modifications that range from simple methylation to complex modifications involving multiple chemical groups (El Yacoubi et al. 2012; Ontiveros et al. 2019). Notably, defects in tRNA modification have emerged as the cause of diverse neurological and neurodevelopmental disorders, thereby highlighting the critical role of tRNA modification in human health and physiology (Angelova et al. 2018; Ramos and Fu 2018). In particular, the brain appears to be exquisitely sensitive to any perturbation in translation efficiency and fidelity brought about by defects in tRNA modifications, as evidenced from the numerous cognitive disorders linked to tRNA modification (Abbasi-Moheb et al. 2012; Alazami et al. 2013; Blanco et al. 2014; de Brouwer et al. 2018; El-Hattab et al. 2016; Khan et al. 2012; Komara et al. 2015; Martinez et al. 2012; Monies et al. 2019; Ramos et al. 2019; Shaheen et al. 2015; Shaheen et al. 2016a; Shaheen et al. 2016b; Shaheen et al. 2019a; Shaheen et al. 2019b).

The methylation of the nucleotide base or sugar by *S*-adenosyl-methionine (SAM)-dependent methyltransferases represents one of the most common post-transcriptional modifications in tRNA (Ayadi et al. 2019; Hori 2014). One of the very first tRNA methyltransferase enzymes to be discovered is the tRNA methyltransferase 1 (Trm1p) enzyme from yeast *Saccharomyces cerevisiae* (Hopper et al. 1982). *S. cerevisiae* Trm1p is imported into the nucleus and mitochondria, where it catalyzes the methylation of a specific guanosine residue at position 26 in numerous tRNAs to yield the N2,N2-dimethylguanosine (m2,2G) modification (Ellis et al. 1987, 1989; Ellis et al. 1986). While *S. cerevisiae* strains lacking Trm1 are viable, Trm1-null strains display temperature sensitivity and defects in tRNA stability that are further exacerbated by combined deletion of either the Trm4p tRNA methyltransferase or the La RNA binding protein (Copela et al. 2006; Dewe et al. 2012). Furthermore, studies in *Schizosaccahromyces pombe* have identified particular tRNAs that are dependent upon Trm1p and/or La for proper folding, aminoacylation and accumulation (Vakiloroayaei et al. 2017). These observations highlight a key role for Trm1p-catalyzed m2,2G modifications in the stabilization of tRNA structure and function.

Two human homologs of yeast Trm1p have been identified by sequence homology that are encoded by the *TRMT1* and *TRMT1L* genes (Buckland et al. 1996; Liu and Straby 2000; Vauti et al. 2007). While the substrates of TRMT1L remain to be discovered, TRMT1 has been demonstrated to be responsible for the majority of m2,2G modifications in the tRNA of human cells (Dewe et al. 2017). Notably, exome sequencing studies have implicated frameshift mutations in *TRMT1* as the cause for certain forms of autosomal-recessive intellectual disability (ID) disorders (Blaesius et al. 2018; Davarniya et al. 2015; Monies et al. 2017; Najmabadi et al. 2011). The ID-associated TRMT1 mutants have been shown to be defective in tRNA binding and enzymatic activity (Dewe et al. 2017). While frameshift mutations in *TRMT1* have been identified in the human population, *TRMT1* missense alleles that help elucidate the functional consequences of tRNA modification deficiency remain to be found. Moreover, the extent to which m2,2G modifications are impacted in patient cells by the TRMT1 mutations is unknown.

Here, we describe a TRMT1 missense variant in an individual presenting with ID disorder and epilepsy. Most significantly, we find that lymphoblastoid cells prepared from the patient exhibit a severe deficit in m2,2G modifications in tRNAs. Moreover, we find that the mutant TRMT1 protein retains the ability to bind tRNA but is unable to rescue formation of m2,2G in TRMT1-deficient cell lines. Our results uncover a single amino acid substitution in TRMT1 that severely impairs tRNA modification function and provide evidence that the neurodevelopmental and cognitive defects caused by *TRMT1* mutations are linked to loss of m2,2G modification.

## Results

### Identification of a novel TRMT1 mutation implicated in ID

We have previously described a “genomics first” approach to patients with ID (Anazi et al. 2017). A high yield of this approach was noted where a likely causal variant was identified in the majority of the >330 patients in that cohort. One of the reported variants in this cohort is a novel missense variant in the *TRMT1* gene (NM_001136035.3 (TRMT1):c.967C>T, p.Arg323Cys) (Figure 1A). Subsequent Sanger sequencing confirmed the homozygous nature of the mutation in the ID-affected index individual (Figure 1B, 13DG1615). This female patient is currently 8 years old with ID, epilepsy and strabismus (Figure 1C). She was conceived via artificial insemination and pregnancy was uneventful. She was delivered at term vaginally but meconium aspiration resulted in respiratory distress and a short admission to neonatal intensive care for 5 days. Her motor development was normal but cognitive development was slow. Her IQ was 73 at the age of 6.5 years (Vineland scale), and she is currently attending special schooling with poor performance. Her medical history is notable for epilepsy that is partially controlled with medications and hearing loss (profound on the right and limited to peripheral auditory involvement on the left). Physical examination revealed normal brain MRI, head size, height and weight. However, she displayed subtle dysmorphism in the form of strabismus, epicanthus and smooth philtrum. The variant NM_001136035.3 (TRMT1):c.967C>T, p.Arg323Cys is completely absent in >4,000 Saudi exomes and is present at a very low frequency in gnomAD (4 heterozygotes, MAF 0.0000159). Moreover, the variant has a consistently deleterious prediction by *in silico* tools DANN, LRT, MutationAssessor, MutationTaster, PROVEAN, FATHMM-MKL and SIFT. These results identify the first missense mutation in the *TRMT1* gene that is linked to a neurodevelopmental disorder.

**Figure 1.**
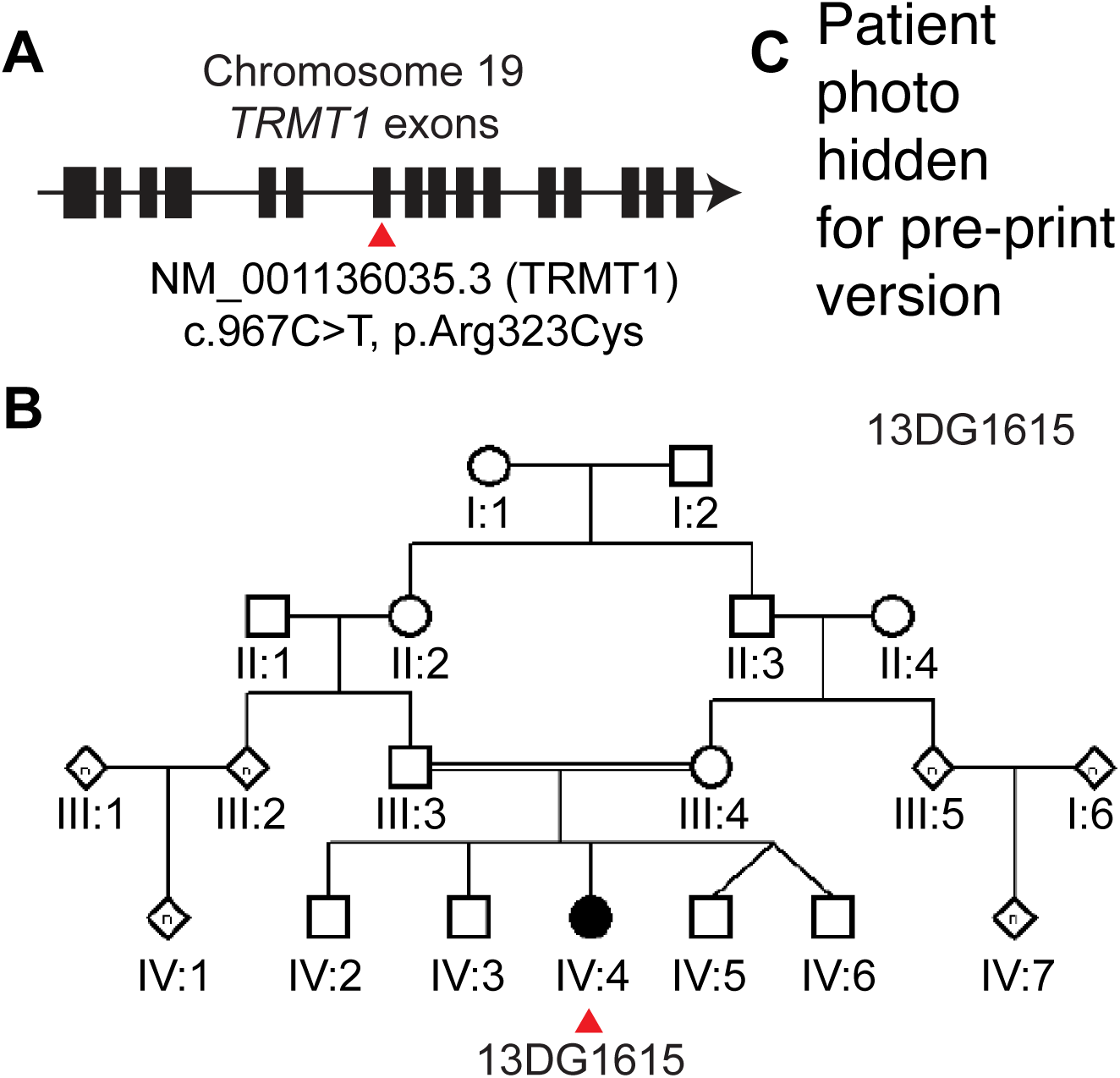
Characterization of a missense mutation in TRMT1 linked to ID. (A) Exon organization of the TRMT1 locus with the location of the single C>T point mutation highlighted in red. (B) Pedigree of the family harboring the missense mutation in the TRMT1 gene and the patient that is homozygous for the mutation. (C) Patient 13DG1615 who is homozygous for the TRMT1 missense mutation who exhibits ID, epilepsy and subtle facial dysmorphism.

### The R323C mutation is predicted to alter TRMT1 folding

The ID-associated variant in the *TRMT1* gene mutates amino acid reside 323 in the SAM-methyltransferase domain of TRMT1 from a positively charged arginine to a non-polar cysteine residue (R323C) (Figure 2A). Based upon protein sequence alignment, the R323 residue of human TRMT1 is absolutely conserved in Trm1 homologs from the Archaea to mammals (Figure 2B). Using the crystal structure of Trm1 from the Archaean *Pyrococcus furiosus* (Ihsanawati et al. 2008), we found that the homologous arginine residue is located within the methyltransferase domain near the putative tRNA binding pocket within a four-stranded beta sheet (Figure 2C). Interestingly, the R323 residue is buried into the interior of Archaeal Trm1 rather than on the surface, unlike most charged side chains which are on the surface of proteins.

**Figure 2.**
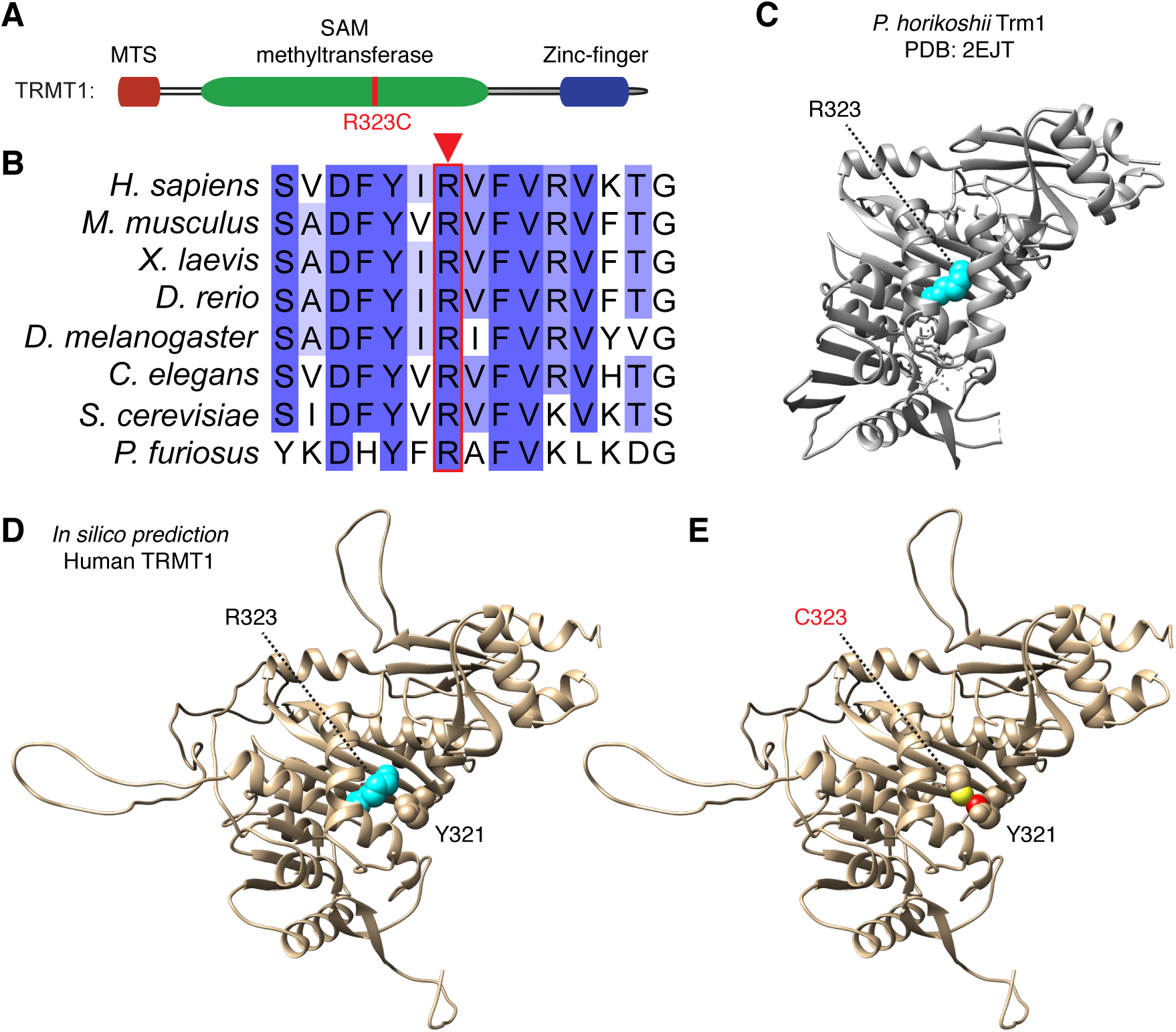
The ID-associated R323C mutation is located at a conserved position in TRMT1 and predicted to perturb protein structure. (A) Schematic of human TRMT1 with protein domains denoted; MTS (mitochondrial targeting signal), SAM-methyl-transferase, and zing-finger motif. The location of the R323C mutation is denoted in red. (B) Protein sequence alignments of TRMT1 from human to Archaea. The R323 residue is boxed in red. (C) The structure of Pyrococcus horikoshi Trm1 (PDB: 2EJT, (Ihsanawati et al. 2008)). The location of the homologous R323 residue is highlighted in cyan. (D) Predicted structure of wildtype TRMT1 based upon in silico template-based tertiary structure prediction (Kallberg et al. 2012). The side chains of the mutated R323 and neighboring tyrosine 321 (Y321) residues are denoted in space-filling form colored cyan and beige, respectively. (E) Predicted structure of TRMT1-R323C with the R323C mutation denoted. Atoms in the side chain of R323C and Y321 that exhibit steric overlap and clash are denoted in yellow and red, respectively.

To gain insight into the potential effects of the R323C mutation on human TRMT1, we generated a predicted tertiary structure of human TRMT1 using an *in silico* template-based algorithm (Kallberg et al. 2012). Based upon this hypothetical structure, TRMT1 is predicted to fold into two distinct domains coinciding with the SAM-methyltransferase domain and the C-terminal CCCH-type Zinc finger motif (Figure 2D, TRMT1 methyltransferase domain). Notably, modeling of the R323C mutation using the most favored rotamer conformation would predict a steric clash with a conserved tyrosine side chain present at position 321 (Figure 2E). Thus, the R323C mutation is likely to be deleterious by perturbing the core packing of the methyltransferase domain of TRMT1 that is responsible for SAM binding, tRNA interaction and catalysis.

### Human patient cells homozygous for the R323C mutation exhibit a severe deficit in m2,2G modification in tRNA

To examine the molecular effects of the R323C mutation, we generated a lymphoblastoid cell line (LCL) from the affected human patient harboring the homozygous missense mutation in the *TRMT1* gene (referred to as R323C-LCL). The R323C-LCL was compared to control lymphoblasts generated from ethnically matched, healthy, unrelated individuals (WT-LCLs). We directly measured and compared the levels of more than 20 different tRNA modifications in the patient R323-LCL versus WT-LCLs through quantitative mass spectrometry of modified ribonucleosides derived from cellular RNA (Cai et al. 2015; Dewe et al. 2017). Strikingly, the m2,2G modification exhibited a 32-fold decrease in the R323-LCL compared to WT-LCL (Figure 3A). No other modification displayed a significant change between the WT versus R323-LCLs.

**Figure 3.**
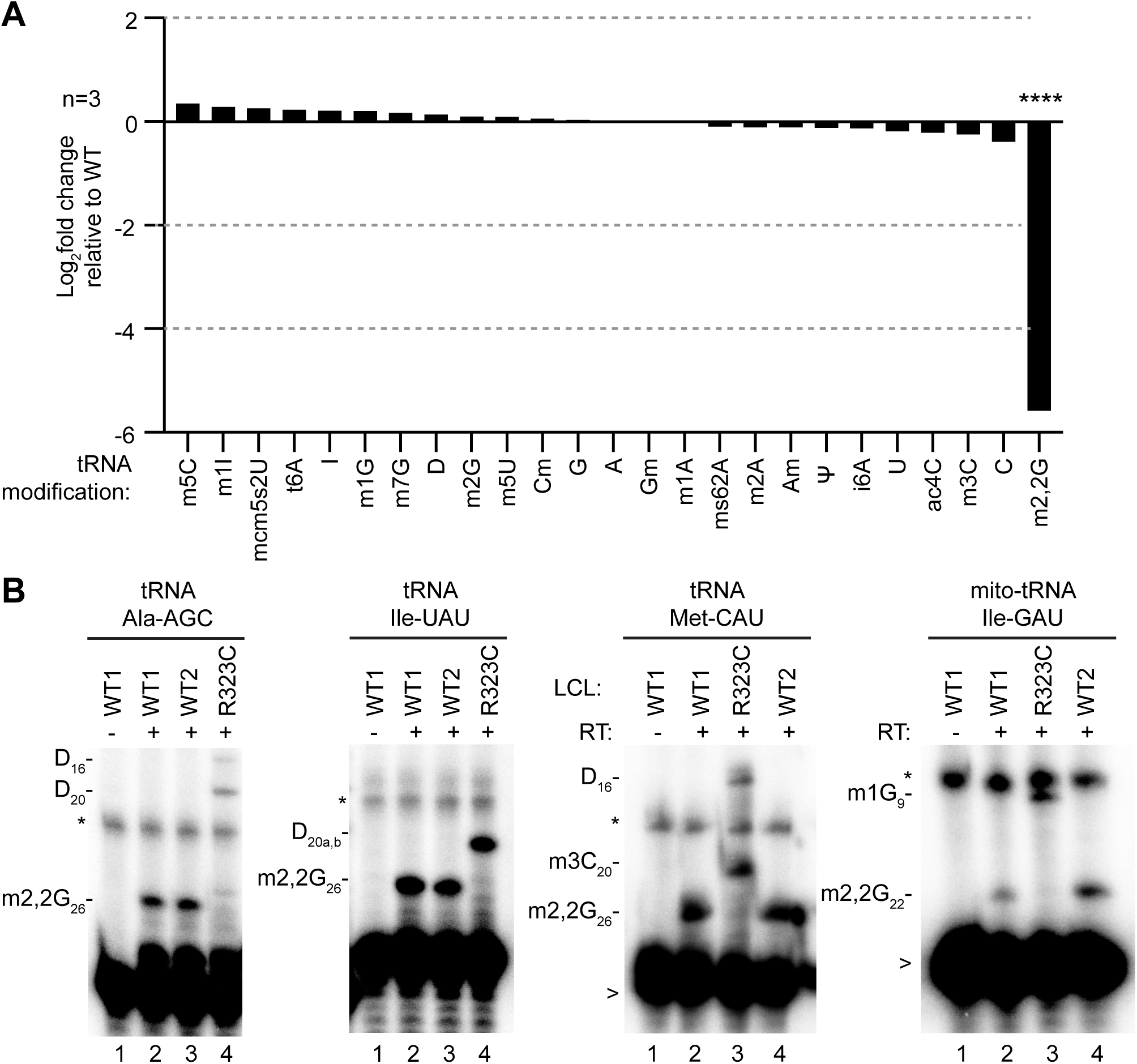
The ID-affected individual expressing only TRMT1-R323C exhibits a severe reduction in m2,2,G modification in tRNA. (A) Comparison of tRNA modification levels between R323C versus WT-LCLs. Nucleosides from digested tRNA samples were analyzed by LC-MS. Y-axis represents the log2-fold change in the levels of the indicated tRNA modification between the R323C patient and WT individual. Samples were measured in triplicate. ****, P < 0.0001 (B) Schematic of primer extension assay to monitor m2,2G in tRNAs. RT, reverse transcriptase. (C) Representative gels of primer extension assays to monitor the presence of m2,2G in tRNA from the indicated LCLs. >, labeled oligonucleotide used for primer extension; D, dihydrouridine; m3C, 3-methylcytosine; m1G, 1-methylguanosine; *, background signal.

To validate the perturbation of m2,2G modification in cellular tRNAs, we used a primer extension assay that detects RNA modification status at nucleotide resolution. In this assay, the presence of m2,2G leads to a block of reverse transcriptase (RT) at position 26 of tRNA while a decrease in m2,2G allows for read-through and an extended product up to a subsequent RT-blocking modification. We selected three nuclear-encoded tRNAs that we have previously shown to contain m2,2G (Dewe et al. 2017), along with mitochondrial tRNA-Ile-GAU, which is the only known mammalian mitochondrial-encoded tRNA to contain m2,2G (Clark et al. 2016; Dewe et al. 2017; Suzuki and Suzuki 2014). In the absence of RT, only background bands were detected in reactions containing the probe and total cellular RNA from the wildtype LCL (Figure 3B, lane 1 for all tRNAs, background bands denoted by *). Addition of RT led to the appearance of an extension product up to the m2,2G modification at the expected position in both nuclear- and mitochondrial-encoded tRNAs in both WT-LCLs (Figure 3B, lanes 2 and 3 for tRNA-Ala-AGC and Ile-UAU or lanes 2 and 4 for tRNA-Met-CAU and mito-tRNA-Ile-UAU). In contrast, the RT block at position 26 was absent in the nuclear- and mitochondrial-encoded tRNA of the R323C-LCLs (Figure 3B, lane 4 for tRNA-Ala-AGC and Ile-UAU or lane 3 for tRNA-Met-CAU and mito-tRNA-Ile-UAU). Loss of m2,2G modification in the tRNAs allowed for readthrough and extension to the next RT-blocking modification. Thus, LCLs from the ID-affected patient with the TRMT1 R323C mutation exhibit a severe defect in m2,2G formation in cellular tRNAs.

### TRMT1 R323C mutant retains the ability to bind RNA but is unable to rescue m2,2G formation in TRMT1-deficient cells

To elucidate the molecular defects associated with the TRMT1-R323C mutant, we investigated the interaction between TRMT1 and tRNAs. As previously shown, human TRMT1 displays a stable interaction with substrate tRNAs that are targets for m2,2G modification (Dewe et al. 2017). Using this system, we expressed a FLAG-tagged version of TRMT1 variants in 293T human embryonic cells followed by affinity purification and analysis of copurifying RNAs. The expressed proteins represent either: 1) wildtype TRMT1, 2) the R323C mutant, and 3) Y445fs, a previously described, ID-associated TRMT1 mutant that lacks RNA binding due to the truncation of the RNA recognition motif (Figure 4A). Immunoblotting confirmed the expression and purification of each TRMT1 variant on anti-FLAG resin (Fig. 4B). In the control purification from vector-transfected cells, we detected only background contaminating 5.8S and 5S rRNAs (Figure 4C, lane 5). In contrast, the purification of WT-TRMT1 resulted in the considerable enrichment of tRNAs along with rRNAs as we have previously shown (Figure 4C, lane 6) (Dewe et al. 2017). Interestingly, we found that similar levels of tRNA were enriched with either TRMT1-WT or TRMT1-R323C mutant (Figure 4C, compare lanes 6 and 7). As expected, the Y445fs mutant exhibited only background RNA signal indicative of defective tRNA binding (Figure 4C, lane 8). Thus, the TRMT1-R323C mutant differs from other ID-associated TRMT1 variants by retaining the ability to bind tRNA.

**Figure 4.**
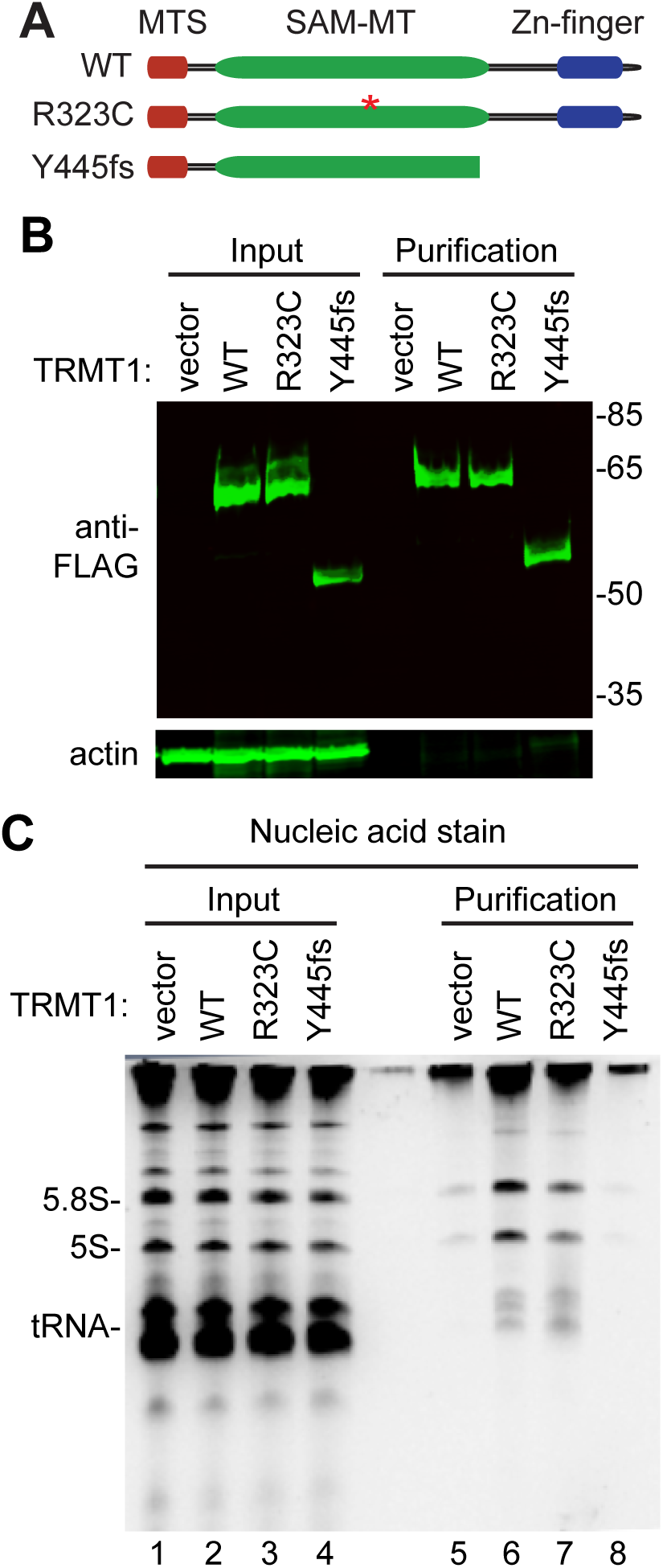
The TRMT1-R323C mutant retains RNA binding but is impaired in reconstitution of tRNA modification activity. (A) Schematic of TRMT1 domains and variants. WT, wildtype; R323C, ID-associated point mutant; Y445fs, TRMT1 variant encoded by an ID-causing frameshift mutation. (B) Immunoblot of whole cell extracts prepared from each human cell line transfected with the indicated constructs. Molecular weight in kiloDalton is denoted on the right. (C) Nucleic acid stain of RNAs extracted from the indicated input or purified samples after denaturing PAGE. The migration pattern of tRNAs, 5.8S and 5S rRNA is denoted.

We next used a previously-described TRMT1-knock out (KO) cell line derived from 293T human cells to dissect the effects of the R323C mutation on TRMT1 function (Dewe et al. 2017). This human 293T cell line lacks TRMT1 expression resulting in the near complete loss of m2,2G modifications in tRNA and the absence of m2,2G modifications in all tested tRNAs. Using transient transfection of mammalian constructs, we expressed either WT-TRMT1 or the R323 variant in the WT or TRMT1-KO 293T cell lines (Figure 5A). We then accessed for rescue of m2,2G formation in tRNA-Ala-AGC using the primer extension assay described above. As expected, non-transfected or vector-transfected WT 293T cells exhibited an RT block at position 26 of tRNA-Ala-AGC indicative of the m2,2G modification (Figure 5B and C, lanes 1 and 2). No readthrough product was detected for either tRNA in WT 293T cells suggesting that nearly all endogenous tRNA-Ala-AGC is modified with m2,2G. Consistent with this observation, increased expression of TRMT1 in WT 239T cells had no noticeable effect on m2,2G modification in tRNA-Ala-AGC (Figure 5B, lane 3). Intriguingly though, over-expression of the TRMT1-R323C mutant in WT 293T cells led to increased readthrough product indicative of decreased m2,2G modification (Figure 5B, lane 4, quantified in 5C). The increase in read-through product suggests that TRMT1-R323C could have a dominant-negative effect on m2,2G modification when over-expressed in the presence of WT-TRMT1.

**Figure 5.**
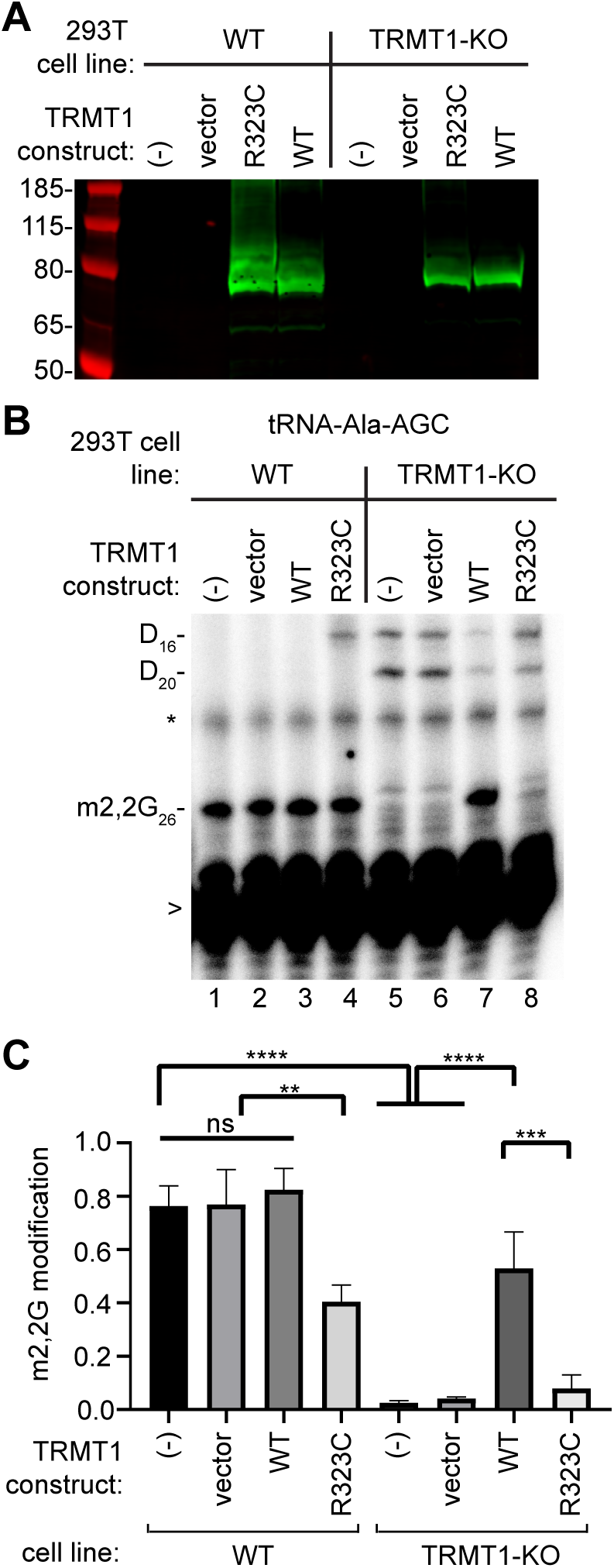
The TRMT1-R323C mutant is impaired in reconstitution of tRNA modification activity in human cells. (A) Immunoblot of whole cell extracts prepared from each human cell line transfected with the indicated FLAG-expression constructs. Blot was probed with anti-FLAG antibody. (B) Representative primer extension assay to monitor the presence of m2,2G in tRNA-Ala-AGC from cell lines transfected with the indicated constructs. >, labeled oligonucleotide used for primer extension; m2,2G, dimethylguanosine; D, dihydrouridine; * background signal. (C) Quantification of m2,2G modification levels in tRNA-Ala-AGC. Primer extensions were performed three times and error bars represent the standard error of the mean. Comparison were performed using one-way ANOVA. *, P < 0.05; **, P < 0.01; ***, P < 0.001; ****, P < 0.0001.

We next tested the TRMT1-R323C mutant for the ability to rescue m2,2G formation in the TRMT1-KO cell line. As expected, the m2,2G modification was absent in tRNA-Ala-AGC isolated from the non-transfected or vector-transfected TRMT1-KO cell line leading to readthrough to the next RT block (Figure 5B, lanes 5 and 6, quantified in 5C). Re-expression of TRMT1-WT in the TRMT1-KO cell line was able to restore m2,2G formation (Figure 5B, lane 7). Due to variable TRMT1 expression caused by incomplete transfection efficiency, the level of m2,2G modification was increased but not completely rescued to the level of the WT cell line. Notably, the TRMT1-R323 mutant displayed greatly reduced ability to reconstitute m2,2G formation in the TRMT1-KO cell line (Figure 5B, lane 8, quantified in 5C). These results indicate that even though TRMT1-R323C can retain binding to tRNAs, it is compromised in its capacity to generate m2,2G in cellular tRNA. Thus, the TRMT1-R323C alteration appears to be a loss-of-function mutation, consistent with the severe deficiency in m2,2G modification in the tRNAs of the affected patient with the R323C mutation.

## Discussion

Here, we characterize a missense mutation in *TRMT1* that causes a cognitive disorder characterized by ID and epilepsy. We further show that cells isolated from the patient with the TRMT1 R323C mutation exhibit a severe deficiency in m2,2G modification within their tRNAs. The TRMT1-R323C mutant retains the ability to bind tRNA but is unable to rescue formation of tRNA modification. These results identify the first missense mutation in the TRMT1 mutation that causes ID and provide the first demonstration that m2,2G levels are impacted in individuals with a pathogenic TRMT1 mutation.

Previously characterized TRMT1-ID mutations result in translation frameshifting that lead to truncation of the carboxyl-terminal zinc finger motif. These mutants have been shown to be defective in RNA binding and reconstitution of methyltransferase activity *in vivo* (Dewe et al. 2017). Unlike the frameshift mutants, the R323C mutant is still able to efficiently interact with tRNA substrates (Figure 4). The retention of tRNA binding by the R323C mutant is consistent with the location of the R323 residue on the interior of TRMT1 within the methyltransferase motif and outside of the putative zinc finger motif that interacts with tRNA. While the R323C mutant can still bind RNA, it exhibits a severe defect in the reconstitution of m2,2G formation in TRMT1-KO human cells. Due to the location of the R323 residue in the core of the TRMT1 methyltransferase domain, the R323C mutation could severely distort the tertiary structure of TRMT1 leading to defects in substrate orientation, SAM binding and/or catalysis.

The m2,2G modification has been predicted to prevent the folding of certain nuclear-encoded tRNAs into alternative conformers found in mitochondrial tRNAs (Steinberg and Cedergren 1995). Moreover, previous studies in *S. cerevisiae* have found that certain tRNA isoacceptors exhibit decreased accumulation in the absence of Trm1, suggesting that the m2,2G modification plays a role in the proper folding of certain tRNAs (Dewe et al. 2012; Vakiloroayaei et al. 2017). Interestingly, we have found that TRMT1-deficient human cells exhibit similar steady-state levels of all tested nuclear- and mitochondrial-encoded tRNAs (Dewe et al. 2017). However, it could be possible that the m2,2G modification is more important for tRNA stability and accumulation for certain human tRNA isodecoders, akin to the situation in *S. cerevisiae*. Thus, it will be interesting to use global profiling approaches to measure tRNA levels to determine if loss of m2,2G impacts some tRNAs more than others. This will allow us to hone in on certain tRNAs that might be more dependent upon the m2,2G modification for processing and stability.

The m2,2G modification could also play a role in proper tRNA interaction with the ribosome to ensure efficient translation. Indeed, studies in yeast S. cerevisiae have found that *Trm1* deletion mutants display alterations in ribosome profiles indicative of translation aberrations (Chou et al. 2017). Moreover, studies in human cells have found that global translation is reduced upon ablation of TRMT1 (Dewe et al. 2017). The alterations in translation are correlated with perturbations in ROS homeostasis and heightened sensitivity to oxidative stress. Future studies using ribosome profiling in patient cells could provide insight into the biological pathways that are perturbed upon loss of the m2,2G modification that contribute to the spectrum of neurodevelopmental phenotypes exhibited by individuals with TRMT1 mutations.

## Materials and Methods

### Human subjects

Evaluation of the affected individual and family members by a board-certified clinical geneticist included obtaining medical and family histories, clinical examination, neuroimaging and clinical laboratory investigations. After obtaining a written informed consent for enrollment in an IRB-approved project (KFSHRC RAC#2070023), venous blood was collected in EDTA and sodium heparin tubes for DNA extraction and establishment of lymphoblastoid cell lines. All studies abide by the Declaration of Helsinki principles.

### *In silico* analysis of TRMT1 structure

The predicted tertiary structure of human TRMT1 (NP_001129507.1) was determined using RaptorX software (http://raptorx.uchicago.edu). Structural visualization, editing, and comparison was performed using UCSF Chimera software (Pettersen et al. 2004). The arginine-323 to methionine mutation was modeled by selecting the most probable rotamer conformation according to the Richardson backbone-independent rotamer library (Lovell et al. 2000). The predicted effects of the R323C mutation on TRMT1 structure was performed using DynaMut (Rodrigues et al. 2018).

### Plasmid constructs

The pcDNA3.1-FLAG-TRMT1-WT and TRMT1-Q415fs expression plasmids have been described previously (Dewe et al. 2017). The pcDNA3.1-FLAG-TRMT1-R323C expression construct was generated by DpnI site-directed mutagenesis. All plasmid constructs have been verified by Sanger sequencing.

### Tissue cell culture

LCLs were grown to confluency and passaged in RPMI 1640 media (ThermoFisher) containing 15% Fetal Bovine Serum, 2 mM L-alanyl–L-glutamine (GlutaMax; Gibco), and 1x penicillin/streptomycin at 37°C with 5% CO_2_. The 293T control-WT and TRMT1-KO cells lines were previously described (Dewe et al. 2017). 293T cells lines were cultured in Dulbecco’s Minimal Essential Medium (DMEM) supplemented with 10% fetal bovine serum (FBS), 1x Glutamax (Gibco), 1x penicillin and streptomycin (ThermoFisher) at 37°C with 5% CO_2_. Cells were passaged every three days with 0.25% Trypsin. For TRMT1 re-expression experiments, pcDNA3.1 vector or pcDNA3.1-TRMT1-expression constructs were introduced into the control-WT or TRMT1-KO1 cell lines using calcium phosphate transient transfection as previously described (Dewe et al. 2017; Kingston et al. 2003). Cells were harvested 48 hours post-transfection for subsequent analysis.

### RNA analysis

RNA was extracted using TRIzol LS reagent (Invitrogen). RNAs were diluted into formamide load buffer, heated to 95°C for 3 minutes, and fractionated on a 10% polyacrylamide, Tris-Borate-EDTA (TBE) gel containing 7M urea. Sybr Gold nucleic acid staining (Invitrogen) was conducted to identify the RNA pattern. For primer extension analysis, 1.5 μg of total RNA was pre-annealed with 5’-^32^P-labeled oligonucleotide and 5x hybridization buffer (250 mM Tris, pH 8.5, and 300 mM NaCl) in a total volume of 7 μl. The mixture was heated at 95°C for 3 min followed by slow cooling to 42°C. An equal amount of extension mix consisting of avian myeloblastosis virus reverse transcriptase (Promega), 5x AMV buffer and 40 μM dNTPs was added. The mixture was then incubated at 42°C for 1 hour and loaded on 15% 7M urea denaturing polyacrylamide gel. Gels were exposed on a phosphor screen (GE Healthcare) and scanned on a Bio-Rad personal molecular followed by analysis using NIH ImageJ software. Primer extension oligonucleotide sequences were previously described (Dewe et al. 2017).

### Liquid chromatography-mass spectrometry

Total RNA from three independent flasks of wildtype and TRMT1-R323C LCLs was isolated using Trizol RNA extraction. Total RNA (20 μg) was digested with nucleoside digestion mix (NEB #M0649) at 37°C overnight. LC-MS analysis of digested nucleosides was analyzed as described previously (Cai et al. 2015; Dewe et al. 2017). Briefly, ribonucleosides were separated using a Hypersil GOLD™ C18 Selectivity Column (Thermo Scientific) followed by nucleoside analysis using a Q Exactive Plus Hybrid Quadrupole-Orbitrap. The modification difference ratio was calculated using the m/z intensity values of each modified nucleoside between WT and R323C-LCL mutants following normalization to the sum of intensity values for the canonical nucleosides; A, U, G and C.

### Protein purification and analysis

Cellular extract was prepared as previously described (Dewe et al. 2017). Briefly, human 293T cells were harvested by trypsinization and washed once with PBS. The cell pellet was resuspended in 500 μL hypotonic lysis buffer (20mM HEPES, pH 7.9; 2 mM MgCl_2_; 0.2 mM EGTA, 10% glycerol, 0.1 mM PMSF, 1 mM DTT), incubated on ice for 5 minutes and subjected to three freeze-thaw cycles in liquid nitrogen and 37°C. NaCl was added to extracts till the final concentration reached 400 mM. After centrifugation at 14,000xg for 15 minutes at 4°C, an equal amount of hypotonic lysis buffer with 0.2% NP-40 was added to 500 μL of soluble cellular extract. FLAG-tagged proteins were purified by incubating whole cell lysates from the transfected cell lines with 20 μL of DYKDDDDK-Tag Monoclonal Antibody Magnetic Microbead (Syd Labs) for three hours at 4 °C. Magnetic resin was washed three times in hypotonic lysis buffer with 200 mM NaCl.

For protein immunoblotting, cell extracts and purified protein samples were boiled at 95°C for 5 minutes followed by fractionation on NuPAGE Bis-Tris polyacrylamide gels (Thermo Scientific). Separated proteins were transferred to Immobilon FL polyvinylidene difluoride (PVDF) membrane (Millipore) for immunoblotting. Membrane was blocked by Odyssey blocking buffer for 1 hour at room temperature followed by immunoblotting with the following antibodies: anti-FLAG epitope tag (L00018; Sigma) and actin (L00003; EMD Millipore). Proteins were detected using a 1:10000 dilution of fluorescent IRDye 800CW goat anti-mouse IgG (925-32210; Thermofisher). Immunoblots were scanned using direct infrared fluorescence via the Odyssey system (LI-COR Biosciences).

## Acknowledgements

We thank the study family for agreeing to participate. We thank members of the Fu Lab for helpful discussion on the manuscript. This work was supported by the Saudi Human Genome Program, and King Salman Center for Disability Research (F.S.A.), and a University of Rochester Furth Fund Award and National Science Foundation CAREER Award 1552126 to D.F..

On behalf of all authors, the corresponding author states that there is no conflict of interest.

